# Dogme: A nextflow pipeline for reprocessing nanopore RNA and DNA modifications

**DOI:** 10.1101/2025.06.04.657941

**Authors:** Elnaz Abdollahzadeh, Ali Mortazavi

## Abstract

**Motivation:** The Oxford Nanopore Technologies (ONT) platform allows for the direct detection of RNA and DNA modifications from unamplified nucleic acids, which is a significant advantage over other platforms. However, the rapid updates to ONT basecalling models and the evolving landscape of computational tools for modification detection bring about challenges for reproducible and standardized analyses. To address these challenges, we developed Dogme, which is a Nextflow-based workflow that automates the processing of ONT data, including basecalling, alignment, modification detection, and transcript quantification. Dogme automates the reprocessing of ONT POD5 files by integrating basecalling using Dorado, read mapping using minimap2 and subsequent analysis steps such as running modkit. The pipeline supports three major types of ONT sequencing data – direct RNA (dRNA), complementary DNA (cDNA), and genomic DNA (gDNA) – enabling comprehensive analyses across different library preparations. Dogme facilitates detection of diverse RNA modifications supported by Dorado such as N6-methyladenosine (m6A), 5-methylcytosine (m5C), inosine, pseudouridine, 2’-Omethylation (Nm) and DNA methylation, while concurrently quantifying full-length transcript isoforms LR-Kallisto for transcript quantification for dRNA and cDNA.

**Results:** We applied Dogme to three separate mouse C2C12 myoblast replicates using direct RNA sequencing on MinION flow cells. We detected an average of 147,879 m6A, 86,673 m5C, 21,242 inosine, 24,540 pseudouridine, and 83,841 2’- O-methylation sites per replicate with 96,581 m6A, 43,446 m5C, 8,825 inosine, 10,048 pseudouridine, and 30,157 2’-O-methylation sites detected in all three biological replicates. The pipeline produced reproducible modification profiles and transcript expression levels across replicates, demonstrating its utility for integrative long-read transcriptomic and epigenomic analyses.

**Availability:** Dogme is implemented in Nextflow and is freely available under the MIT license at https://github.com/mortazavilab/dogme, with documentation provided for installation and usage.

## Introduction

Long-read sequencing technologies have revolutionized genomics research by enabling the analysis of complex genomic loci (1) as well as full-length transcripts (2) that were previously difficult to characterize with short-read approaches alone. Oxford Nanopore Technologies (ONT) supports the direct detection of native RNA and DNA modifications on unamplified RNA (3) and DNA (4) molecules.

Oxford Nanopore Technologies (ONT) stores the raw electrical signal generated by either the MinION or PromethION flowcells as each RNA or DNA molecule passes through a tiny nanopore into a POD5 file after a fixed number of reads or time, depending on user preferences. Thus each sequencing run typically generates dozens to hundreds of POD5 files, which must be processed through several computational steps such as basecalling, mapping and modification calling to extract biologically meaningful information. While the Minknow user interface automates much of this for an initial run, not all basecalling features, including some of the newest modifications are available in a given release. Furthermore, ONT has been rapidly updating their Dorado basecalling software and models multiple times a year to achieve higher read quality, which can only be leveraged by reprocessing the raw POD5 data followed by repeating downstream analyses. This is a time-consuming process to repeat on a large number of samples that must be analyzed uniformly in large consortia such as ENCODE (5) and IGVF (6). To address the challenge of uniformly reprocessing samples generated across a long period of time such as the lifetime of a consortium, we developed Dogme, which is a Nextflow-based workflow that automates the processing of ONT raw signaling data through basecalling, alignment, modification detection, and transcript quantification. This workflow leverages the latest ONT basecalling models while ensuring computational reproducibility.

## Method

Dogme is implemented in Nextflow, which enables scalable and reproducible scientific workflows across multiple computing environments (7). Dogme is specifically designed to handle three distinct types of sequencing data: direct RNA (dRNA) for detecting native RNA modifications, complementary DNA (cDNA) for high-accuracy transcript quantification without modifications, and genomic DNA (gDNA) for epigenetic analysis. All of the information to run Dogme is stored in a single configuration file, which specifies the sample type, the location of the starting POD5 files and all of the output folders, the modifications to be analyzed as well as the reference genomes and transcriptomes. The config file also has a section that specifies cluster-specific settings and queues. A separate file allows users to specify the paths of the pre-installed copies of Dorado (8), Samtools (9), Minimap2 (10), Modkit (11), and Kallisto-bustools (12). The default Dogme workflow (Fig. 1A) starts by downloading the latest Dorado models, then proceeds with base-calling individual POD5 files using Dorado in parallel to produce one BAM file per POD5, which are then merged into a single unmapped BAM file stored in the bams folder. When used to analyze DNA or RNA modifications, this unmapped “modBAM” holds both the sequence information and modification probability scores for each nucleotide position. For dRNA and cDNA, a Fastq file is extracted from the BAM, which is then used to run long-read kallisto and bustools to quantify gene expression using a precomputed kmer index.

**Fig. 1.**
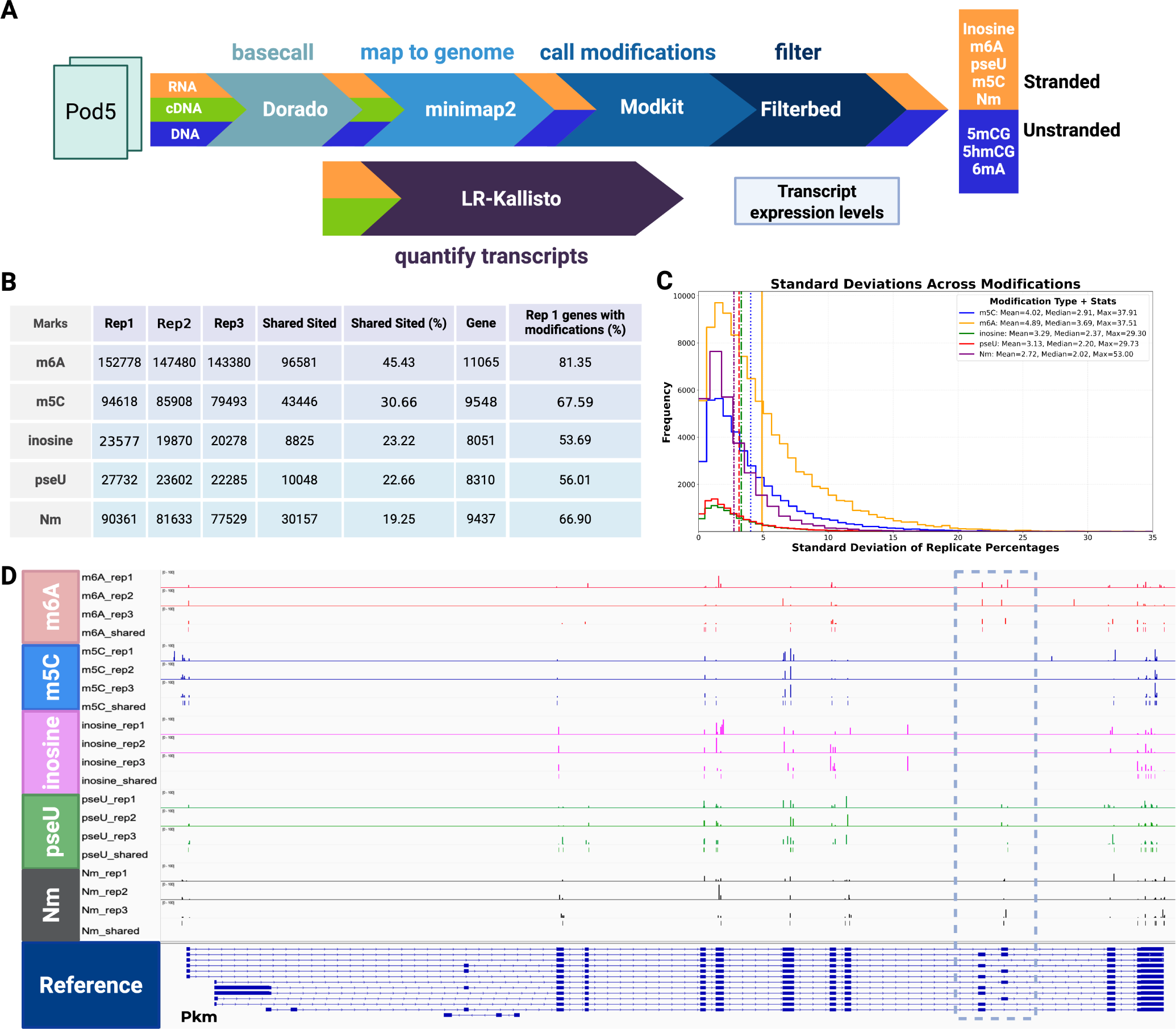
A uniform pipeline for reprocessing Nanopore data. **A**. Dogme workflow for processing dRNA (orange), cDNA (green), and gDNA (blue) Nanopore reads through several steps including basecalling, alignment, modification detection (RNA and gDNA), and expression quantification (dRNA and cDNA). **B**. Table of detected RNA modifications across three C2C12 myoblast replicates with the percentage of expressed genes in rep1 that are modified. **C**. Distribution of standard deviations in modification percentages, showing Poisson-like behavior. **D**. RNA modification profiles for *Pkm* with 5’ on the left, highlighting region-specific patterns of m6A, pseudouridine, m5C, inosine, and Nm for 2’-O-methylation of any of the four bases. The height of the modifications represent the percentage of modifications at that position in each replicate. Sites that are shared between the three replicates are shown in their own tracks. Dashed box shows alternatively spliced exons that have reproducible sites in one exon or the other.

For all three sample types, reads are aligned to the genome using Minimap2. For dRNA and cDNA, Minimap2 uses the splice junctions from the gene annotation GTF file as a guide. Following Minimap2 alignment, the pipeline employs Modkit to extract RNA and DNA modifications from the aligned BAM files. Modkit leverages basecalling probabilities and positional information to identify modified bases either by sampling reads or using a user-defined threshold specified in the Dogme configuration file. Dogme finally applies an adjustable custom filtering step with thresholds of at least 3 supporting reads and a minimum modification frequency of 5 percent to minimize false positives while maintaining sensitivity. Importantly, Dogme is compatible with both stranded and unstranded data and supports analysis of both the plus and minus strands for RNA modifications, ensuring comprehensive coverage of transcriptomic and epigenetic features across all orientations.

It is worth noting that Dorado is a GPU-intensive program, whereas all downstream steps are CPU-heavy. While Dogme is designed to start from running Dorado on POD5 files in the default workflow, users have the option to run Dogme with an alternative workflow to remap reads by skipping Dorado and start from an existing unmapped BAM file.

## Results

We tested Dogme using bulk directRNA sequencing from three biological replicates of mouse C2C12 myoblasts from the ENCODE project, on separate ONT MinION R10.4.1 flow cells sequenced to an average depth of 3.5 million reads per sample (ENCODE accession: ENCSR160HKZ). Reads were mapped on GRCm39 and quantified using GENCODE v36 annotations. We ran Dorado 1.0 on a SLURM HPC cluster with separate queues for NVIDIA A100 GPUs for the Dorado tasks and CPUs for the remaining tasks. Dorado currently supports the detection of five RNA modifications: N6methyladenosine (m6A), 5-methylcytosine (m5C), inosine (A-to-I), pseudouridine (Ψ), and 2’-O-methylation on any base (Nm). These five modifications play crucial roles in regulating gene expression, RNA stability, splicing, and translation, making them essential components of cellular processes and potential biomarkers for disease (13). We processed each sample through Dogme and detected 21,138 to 21,476 expressed transcripts in 13,058 to 13,278 genes per replicate. We identified an average of 147,879 (±3,841) unique m6A sites, 86,673 (±6,197) m5C sites, 21,242 (±1,663) inosine sites, 24,540 (±2,321) Ψ sites, and 83,841 (±5,369) Nm sites per sample on non mitochondrial chromosomes (Fig 1B). Pairwise comparisons of modification levels demonstrated strong concordance between replicates, with consistent sitespecific modification percentages and high reproducibility as evidenced by scatter plot correlations (not shown).

Density distribution analysis revealed that while most sites exhibited relatively low modification levels, a subset showed high modification percentages across all replicates, suggesting biological significance rather than technical artifacts. Standard deviation analysis of modification percentages across all detected sites revealed similar Poisson distributions for all five modification types, further confirming the technical reproducibility of our approach (Fig. 1C).

Examination of the gene Pkm (pyruvate kinase, muscle) illustrated the pipeline’s ability to detect modification patterns with high reproducibility. Across the three replicates, we consistently identified 21-29 m6A sites, 26-31 m5C sites, 27-31 Nm sites, 28-39 Ψ sites, and 18-24 low-frequency inosine sites within this gene. 14 m6A, 23 m5C, 17 Nm, 21 Ψ, and 12 inosine sites were shared across all replicates in Pkm. Interestingly, some of the shared modifications are found in alternatively spliced exons (Fig. 1D). Analysis of the modifications using long-reads could therefore be used to tie modification patterns to specific isoforms. These results demonstrate that the Dogme pipeline enables robust and reproducible detection of RNA modifications from direct RNA sequencing data, providing a foundation for investigating epitranscriptomic dynamics in diverse biological contexts.

## Conclusions

The development and validation of the Dogme pipeline addresses the computational challenges of uniformly reprocessing RNA and DNA modifications to leverage the latest ONT models. Our results from C2C12 myoblasts demonstrate that this approach can consistently identify and quantify diverse RNA modifications across biological replicates with high reproducibility. It is remarkable that we detect each modification in at least half of the expressed genes in a sample (Fig. 1B). Furthermore, the ability to detect these patterns reproducibly across replicates indicates that Dorado can recover biological signals that are not technical noise, a critical requirement for confident epigenomic and epitranscriptomic analysis. While Dogme is not designed to analyze RNA modifications on specific transcripts, there are clear patterns of modifications that are isoform specific in the C2C12 data that would be very interesting to follow up on. Future directions for the Dogme pipeline include expanding modification detection capabilities, integrating additional downstream analysis tools for functional interpretation, adding support for single cell long read cDNA, and adapting the workflow for comparative analysis across different tissues, conditions, or species.

## Acknowledgments

We thank Luis Solano, Jasmine Sakr, and Negar Mojgani for preparing cells and sequencing the samples, Diane Trout for assistance with uploading the C2C12 datasets to the EN-CODE portal, and Mortazavi lab members for discussions.

